# Metastasis of colon cancer requires Dickkopf-2 to generate cancer cells with Paneth cell properties

**DOI:** 10.1101/2024.04.12.589235

**Authors:** Jae Hun Shin, Jooyoung Park, Jaechul Lim, Jaekwang Jeong, Ravi K. Dinesh, Stephen E. Maher, Jeonghyun Kim, Soyeon Park, Jun Young Hong, John Wysolmerski, Jungmin Choi, Alfred L. M. Bothwell

**Affiliations:** Integrative Science and Engineering Division, Underwood International College, Yonsei University, Incheon 21983, Korea; Institute of Advanced Bio-Industry Convergence, Yonsei University, Seoul, Korea; Department of Biomedical Sciences, Korea University College of Medicine, Seoul 02841, Korea; College of Veterinary Medicine, Seoul National University, Seoul 08826, Korea; Internal Medicine, Yale University School of Medicine, New Haven, Connecticut 06520, U.S; Department of Pathology, Stanford University, Stanford, California 94305, U.S; Department of Urology, Yale University School of Medicine, New Haven, Connecticut 06520, U.S.; Department of Systems Biology, College of Life Science and Biotechnology, Yonsei University, Seoul, Korea; Dept. of Pathology, Microbiology and Immunology, University of Nebraska Medical Center, 505 S. 45^th^ Street., Omaha, NE 68198, U.S; Dept. of Immunobiology, Yale University School of Medicine, New Haven, Connecticut 06520, U.S

**Author notes:** Correspondence to Alfred L. M. Bothwell, Dept. of Pathology, Microbiology and Immunology, University of Nebraska Medical Center, 986842 Nebraska Medical Center, Fred & Pamela Buffett Cancer Center, 505 S. 45^th^ Street, Omaha, NE 68198, Jae Hun Shin, Integrative Science and Engineering Division, Underwood International College, Yonsei University, Incheon 21983, South Korea, 82-10-2676-0525. These authors are equally contributed (Jae Hun Shin, Jooyoung Park).

**Keywords:** Metastasis, Colon cancer, DKK2, Paneth cell properties

## Abstract

Metastasis is the leading cause of cancer-related mortality. Paneth cells provide stem cell niche factors in homeostatic conditions, but the underlying mechanisms of cancer stem cell niche development are unclear. Here we report that Dickkopf-2 (DKK2) is essential for the generation of cancer cells with Paneth cell properties during colon cancer metastasis. Splenic injection of *Dkk2*-knockout (KO) cancer organoids into C57BL/6 mice resulted in a significant reduction of liver metastases. Transcriptome analysis showed reduction of Paneth cell markers such as lysozymes in KO organoids. Single cell RNA sequencing analyses of murine metastasized colon cancer cells and patient samples identified the presence of lysozyme positive cells with Paneth cell properties including enhanced glycolysis. Further analyses of transcriptome and chromatin accessibility suggested Hepatocyte nuclear factor 4-alpha (HNF4A) as a downstream target of DKK2. Chromatin immunoprecipitation followed by sequencing analysis revealed that HNF4A binds to the promoter region of *Sox9*, a well-known transcription factor for Paneth cell differentiation. In the liver metastatic foci, DKK2 knockout rescued HNF4A protein levels followed by reduction of lysozyme positive cancer cells. Taken together, DKK2-mediated reduction of HNF4A protein promotes the generation of lysozyme positive cancer cells with Paneth cell properties in the metastasized colon cancers.

## Introduction

Metastasis is the main cause of cancer-related death (1). Mutations in the *Apc* gene in intestinal epithelial cells cause adenoma formation by dys-regulation of Wnt signaling (2). Accumulating oncogenic mutations in *Kras*, *Tp53* and *Smad4* genes facilitate colorectal carcinogenesis and metastasis (3–5). Recent studies have identified that *Lgr5* positive cells are required for metastasis in the murine models of colorectal cancer (6, 7). *Lgr5* positive cells include stem cells, transit amplifying cells and Paneth cells (8). Ablation of *Lgr5* positive cells in the primary tumor in mice restricted metastatic progression of colon cancer cells (6). Time-lapse live imaging analysis of metastatic cancer cells in the murine primary colon tumors revealed that both *Lgr5* positive and negative cancer cells are able to escape from the primary tumors and circulate in the blood stream (7). Importantly, generation or presence of *Lgr5* positive cells is necessary to form metastases suggesting that cancer stem cells and their niche formation upon metastases seeding is a prerequisite of metastatic tumor growth.

The stem cell niche provides so called ‘stem cell niche factors’ in order to regulate proliferation and differentiation of stem cells (9, 10). In the homeostatic condition, Paneth cells function as stem cell niche cells by providing Wnt3, EGF, Notch ligand, and Dll4 to intestinal stem cells (11). Paneth cell-mediated glucose conversion into lactate promotes proliferation of intestinal stem cells that lack glucose metabolism (12). Stem cell niche factors not only maintain stem cells but also generate new stem cells by dedifferentiation from the progenitors when the original stem cells are damaged by intestinal injury or inflammation (13–15). Regeneration of stem cells by niche factors also occurs in the context of colorectal cancer (6, 16, 17). However, the underlying mechanisms of cancer stem cell niche cell development in the primary and metastatic colon tumors are largely unknown.

In normal mucosa, the balance between Wnt and Notch signaling determines the fate of intestinal stem cells differentiating to secretory or absorptive precursors (18–20). *Atoh1* expression in the secretory precursors directs their differentiation to Paneth or goblet cells (21). *Sox9* expression then induces Paneth cell differentiation (22, 23) in the small intestine. Recent findings have identified the presence of Paneth-like cells in the large intestine (24–26). However, the existence of Paneth or Paneth-like cells in colon cancers remains unknown. Using colon cancer organoids carrying mutations in *Apc*, *Kras* and *Tp53* genes, we showed here that *Dkk2* knockout disrupted Lysozyme (LYZ) positive cell formation in colon cancer organoids as well as metastatic nodules *in vivo*. Single cell RNA sequencing analyses for mouse and human metastatic colon cancer samples have identified the existence of LYZ+ cells exhibiting Paneth cell properties including metabolic support for stem cells (27). Mechanistically, DKK2 protein deficiency recovered protein levels of Hepatocyte Nuclear Factor 4 alpha (HNF4A) in colon cancer cells (28). HNF4A binding on the *Sox9* promoter inhibited the formation of LYZ+ cancer cells exhibiting Paneth cell characteristics. The loss of LYZ+ cells by *Dkk2* knockout rescued mice from the liver metastasis of colorectal cancer induced by cancer organoid transplantation. Our findings suggest that DKK2 promotes LYZ+ cell formation exhibiting Paneth cell properties to develop cancer stem cell niches for outgrowth of metastasized colon cancers.

## Results

### DKK2 is indispensable for liver metastasis of colorectal cancer

Metastatic potential of *Lgr5*-expressing cells has been reported in colorectal cancer (29). The ablation of *Lgr5*-expressing cells in primary colon tumors inhibited metastasis whereas the growth of primary tumors quickly recovered by dedifferentiation of non-stem cells into *Lgr5-* expressing stem cells (6). Moreover, *Fumagalli et al.* have shown that *Lgr5*-expressing cells are indispensable for the growth of metastases (7). We have recently reported that DKK2 enhanced *Lgr5* expression in colon cancers through activation of c-Src (28). Enhanced activation of c-Src has been reported in metastatic colon cancers as well (30). Based on these observations, we queried whether DKK2 is required for metastasis of colorectal cancer. We developed colon cancer organoids carrying oncogenic mutations in *Apc*, *Kras* and *Tp53* genes (AKP) (28). Splenic injection of the *Dkk2* knockout AKP (KO) organoids into wild type C57BL/6 mice resulted in significantly less liver metastasis and mortality compared to the control (AKP) organoids (Fig. 1A-D). Quantitative gene expression analyses revealed that the levels of *Dkk2* and *Lgr5* were still decreased in KO-derived liver metastasized cells compared to AKP-derived cells (Fig. 1E). Lgr5 protein expression and c-Src phosphorylation were reduced in KO-derived liver tumors (Fig. 1F and G). The percentage of *Lgr5*-expressing cells with c-Src phosphorylation in KO tumor tissues was decreased about 2-fold compared to AKP tissues (Fig. 1H). These data suggest the necessity of DKK2 in the growth of *Lgr5*-expressing cancer stem cells in liver metastasized nodules originated from the splenic injection of cancer organoids.

**Figure 1.**
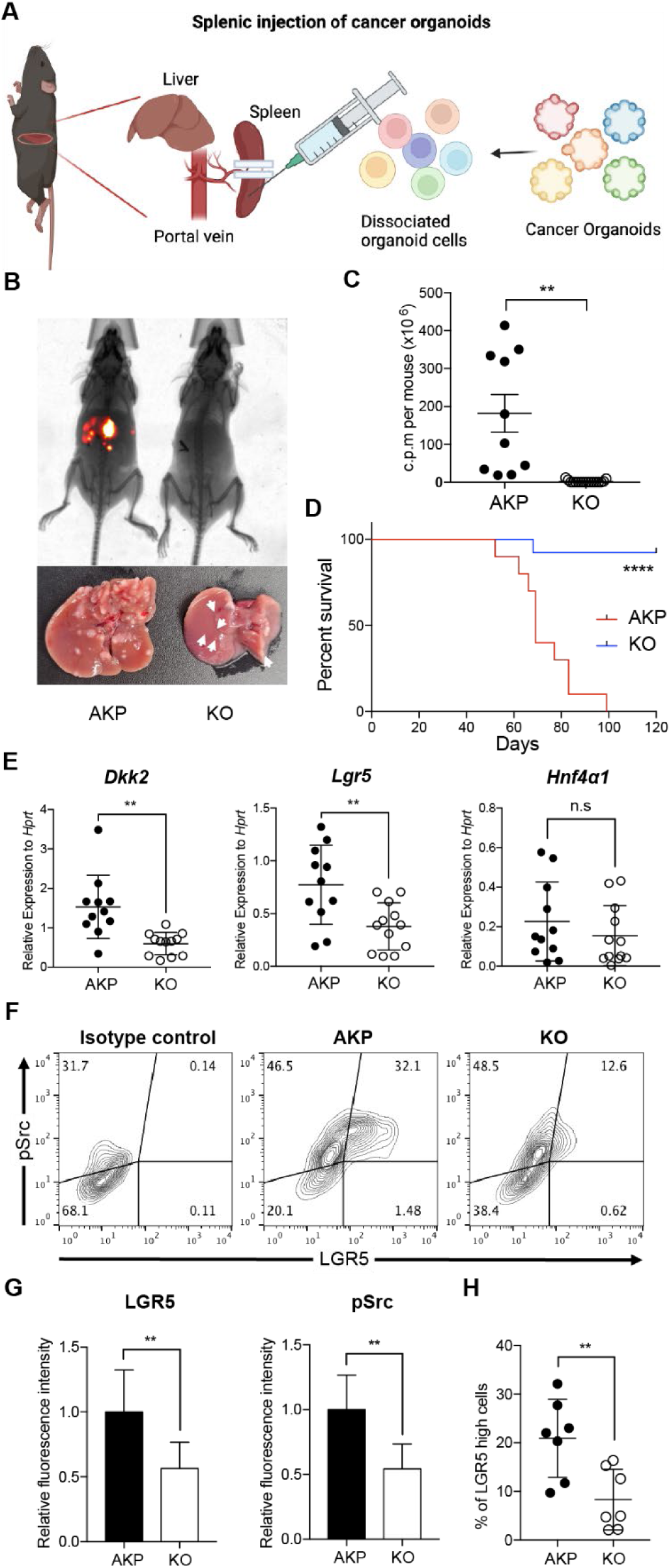
DKK2 is required for Lgr5 positive stem cells-driven liver metastasis of colorectal cancer. **(A)** Isolated cells from AKP tumor organoids expressing the *tdTomato* reporter gene were dissociated and injected into the spleen of wild type C57BL/6 mice. 8 weeks after the injection, metastatic tumor growth was measured by in vivo imaging analysis. **(B)** Representative pictures of *in vivo* imaging analysis. Ctrl: AKP control organoids transduced with scrambled guide RNA, KO: *Dkk2* knockout AKP organoids. **(C-D)** Statistic analysis of liver metastasis (C) and survival (D). Ctrl (*n*=10), KO (*n*=15), *n* represents the number of mice. c.p.m in (C): count per minutes **(E)** Quantitative gene expression analysis of *Dkk2, Lgr5* and *Hnf4*α*1* in liver metastasized colon cancer cells. Ctrl (*n*=11), KO (*n*=12), *n* represents the number of isolated cancer nodules with 5 mice per group. **(F)** Representative flow cytometry analysis data of c-Src phosphorylation (pSrc) and *Lgr5* expression in metastasized colon cancer cells. **(G)** Statistic analysis of *Lgr5* expression and pSrc in (E). The average of mean fluorescence intensity in control samples was set as 1 and relative fluorescence intensity was calculated. **(H)** Statistic analysis of the percentile of Lgr5 high (Lgr5 and pSrc double positive) cells in metastasized colon cancer in (F). Each *symbol* represents an individually isolated cancer nodule. n.s = not significant, **P<0.01, ****P<0.0001; two-tailed Welch’s t-test (C, E, G, H). Error bars indicate mean ± s.d. Log-rank test (D). Results are representative of three independent experiments.

### DKK2 is required for lysozyme positive cell formation in colon cancer organoids

To understand the underlying mechanisms of the DKK2-mediated increase of *Lgr5*- expressing stem cells in liver metastases, we performed RNA sequencing (RNA-seq) analysis on AKP and *Dkk2* knockout (KO) organoids. KO organoids were generated using the CRISPR technique and DKK2 reconstitution was followed using recombinant mouse DKK2 protein (Fig. 2A). Consistent with our previous report using a colitis-induced tumor model, expression of stem cell marker genes including *Lgr5* was reduced in KO organoids (Fig. S1A) (28). In particular, the expression of Paneth cell marker genes such as *Lyz1*, *Lyz2*, and alpha defensins were significantly reduced in KO organoids (Fig. 2B and S1B). Paneth cells are derived from *Lgr5*- expressing stem cells by asymmetric division and maintain their *Lgr5* expression (15). These cells produce stem cell niche factors such as Wnt3A in order to facilitate stem cell proliferation and differentiation into other epithelial lineage cells (11). Paneth cells are also crucial for the glycolysis of intestinal stem cells that lack of glucose metabolism (12). Our bulk RNA-seq data showed that Paneth cell-derived Wnt3a as well as *Fgfr3*, a marker of Paneth cell-induced expansion of stem cells were reduced by *Dkk2* knockout, indicating that the formation of stem cell niche might be disrupted in KO organoids (Fig. S1C) (31). Indeed, formation of Lysozyme positive (LYZ+) cells was disrupted by *Dkk2* knockout during organoid culture (Fig. 2C). LYZ+ cells were detected in AKP organoids from day 2 after plating a single or multiple cells of enzyme-digested organoids whereas those cells were less frequently observed in KO organoids. It correlates to the expression pattern of *Lgr5* in our previous report (28). A significant increase of *Lgr5* expression was observed between day 2 and day 4 and peaked at day 4 to day 6 in AKP organoid culture.

**Figure 2.**
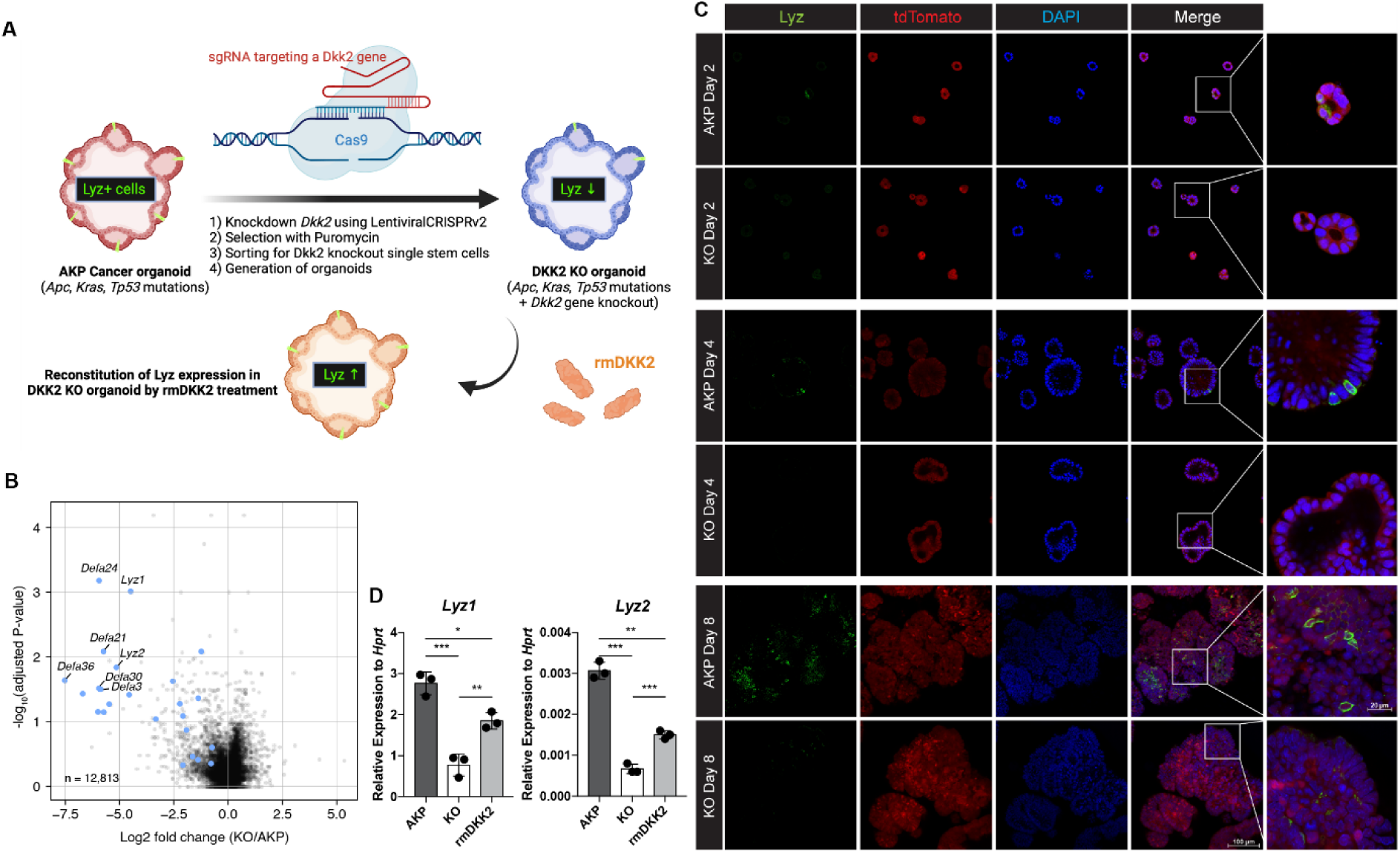
DKK2 is indispensable for the generation of cancer cells with Paneth cell properties in colon cancer organoids. **(A)** A schematic diagram of the generation of KO organoids using CRISPR technique and the reconstitution of DKK2 in organoids by recombinant mouse DKK2 protein (rmDKK2) treatment. Lysozyme (LYZ) expression is highlighted for the followings. **(B)** A volcano plot of RNA-seq analysis comparing KO versus AKP organoids. Paneth cell marker genes are highlighted as blue circles (AKP=3 and KO=5 biological replicates were analyzed). **(C)** Confocal microscopy analysis of LYZ positive cells in AKP or KO organoids in a time-dependent manner using anti-Lysozyme (LYZ) antibody. **(D)** Quantitative real time PCR analyses of *Lyz1* and *Lyz2* in 8 days cultured colon cancer organoids. KO organoids were cultured in the presence of 1 μg/ml of recombinant mouse DKK2 protein. *P<0.05, **P<0.01, ***P<0.001; two-tailed Welch’s t-test. Error bars indicate mean ± s.d. Results are representative of three biological replicates.

*Lgr5* expression was reduced about 3-fold in KO organoids compared to AKP organoids at day 8 (28). Likewise, the markers of Paneth cells, *Lyz1* and *Lyz2* were reduced in KO organoids at day 8 and recombinant mouse DKK2 protein treatment into KO organoids partially rescued it (Fig. 2D). These data indicate that DKK2 is necessary for LYZ+ cell formation in colon cancer organoids that might be required for the cancer stem cell niche formation.

### LYZ+ cancer cells exhibit Paneth cell properties in both mouse and human systems

To investigate the necessity of DKK2 in the formation of LYZ+ cancer cells during colon cancer metastasis, we employed the murine *in vivo* model of liver metastasis shown in Figure 1. In order to analyze transcriptome changes by *Dkk2* knockout in each cell type of metastasized tumors, we performed single-cell RNA sequencing (scRNA-seq) analysis on liver metastasized AKP and KO tumor cells. The Louvain method revealed 4 clusters in our dataset. A cell type was assigned to each cluster on the basis of gene expression patterns and visualized in a Uniform manifold approximation and projection (UMAP) representation (Fig. 3A). Epithelial tumor cells were sub-clustered as 7 clusters and the cluster 6 has been identified as cancer cells with Paneth cell properties based on the expression of Paneth marker genes using the Panglao database (Fig. 3A, S2A-D). Total of 17 cells in cluster 6 were further defined as cancer cells with Paneth cell properties (Paneth+) or goblet cell properties (Goblet+) using the relevant marker genes expression, *Lyz1*, *Muc2* and *Atoh1* (Fig. S2E and S2F). The calculated ratio between Paneth+ cells and Goblet+ cells in KO metastasized tumors was 0.43, reduced as half compared to the ratio in the AKP control, 1 (Fig. S2F). This suggests a reduction of LYZ+ cells in metastasized colon tumors by knockout a *Dkk2* gene.

**Figure 3.**
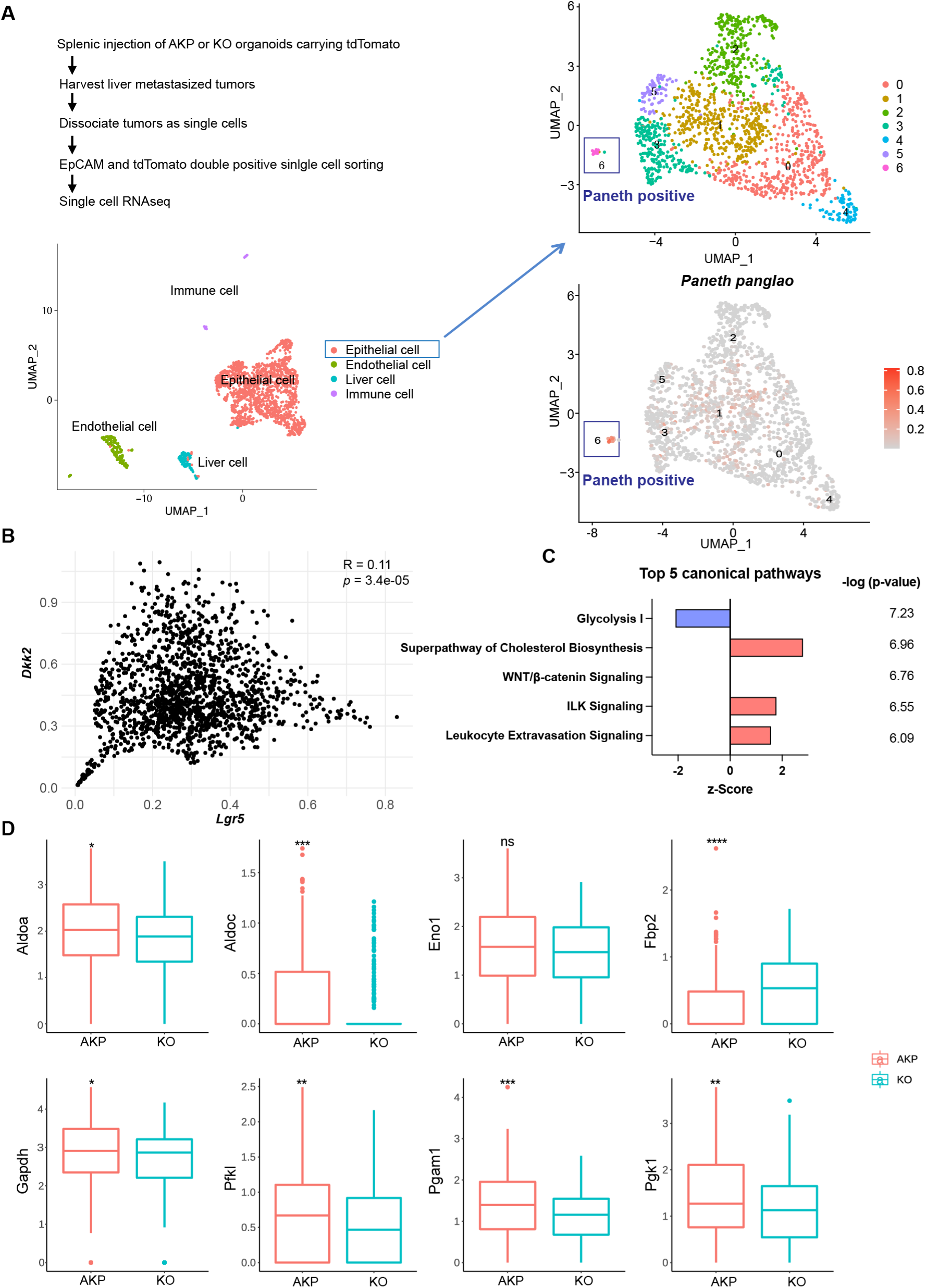
Cancer cells harboring Paneth cell properties were reduced in *Dkk2* knockout metastasized colon cancer tissues in mice. Control AKP or KO colon cancer organoids were transplanted via splenic vein as described in Figure 1. 3 weeks after transplantation, mice were sacrificed to analyze metastatic tumor growth in liver. **(A)** Single cell RNA sequencing analysis (scRNA-seq) of liver metastasized colon cancer tissues. The uniform manifold approximation and projection (UMAP) plot clustered epithelial, endothelial, liver and immune cells in metastasized cancers based on transcriptome analysis. The cancer epithelial cell cluster was sub-clustered to identify cells with Paneth cell properties (cluster 6, Paneth positive). **(B)** The correlation between *Dkk2* and *Lgr5* expression in the cluster of epithelial cells by Pearson r test. **(C)** Ingenuity pathway analysis (IPA)-suggested top 5 canonical pathways of the scRNA-seq data of *Lgr5* positive epithelial cells in KO compared to AKP. z-Scores indicate activation or inhibition of the suggested pathways. The significance values for the pathways are calculated by the right-tailed Fisher’s exact test. **(D)** Box plots show expressions of the genes involved in the glycolysis I pathway in (C). ns = not significant, *<P<0.05, **P<0.01, ***P<0.001, ****P<0.0001; Wilcoxon signed-rank test.

To analyze the consequence of the loss of LYZ+ cells by *Dkk2* knockout in cancer stem cell niche formation, we enriched *Lgr5* positive cells that consist of stem cells, stem cell niche cells and transit amplifying cells in the homeostatic condition (Fig. S3A and S3B) (8). Elevated expression of stem cell marker genes represents cancer stemness in *Lgr5* positive cells compared to *Lgr5* negative cells (Fig. S3C). The UMAP plot of *Lgr5* positive cells displayed 2 clusters (Fig. S3D). Interestingly, expression of Noggin (*Nog*), one of the stem cell niche factors was detected mostly in cluster 1 where only 1 cell was counted in KO (Fig. S3E-S3G). Noggin is an inhibitor of BMP signaling, which restricts stemness of *Lgr5* positive stem cells in the gut (32, 33). Following the reduction of *Nog*, *Bmp4* expression was reversed in KO tumor epithelial cells in the scRNA-seq data (Fig. S3H). This has been confirmed by quantitative gene expression analysis of *Bmp4* in KO organoids (Fig. S3I). The correlative expression between *Dkk2* and the stem cell marker gene, *Lgr5* has been shown in the scRNA-seq data (Fig. 3B). Furthermore, pathways enrichment analysis of the scRNA-seq data using IPA suggested that the glycolysis I pathway was significantly inhibited in KO *Lgr5* positive tumor epithelial cells (Fig. 3C). Expression of glycolysis-related genes such as *Aldoa*, *Aldoc* and *Fbp2* was reduced by *Dkk2* knockout in *Lgr5* positive cells (Fig. 3D) (34). Previous studies have defined that Paneth cells participate in the regulation of energy metabolism in intestinal stem cells (12, 35). Since stem cells are not able to generate lactate, Paneth cells instead process glucose in order to provide lactate to stem cells (12). Therefore, reduction of the glycolysis I pathway in the absence of DKK2 indicates the impaired stem cell niche formation by the loss of LYZ+ cells carrying Paneth cell properties.

Paneth cells constitute the stem cell niche in the small intestine (21). Recent studies have identified the presence of Paneth-like cells in the colon (25, 26). However, the presence of Paneth cells or Paneth-like cells within the tumor microenvironment is elusive. To discern the existence of cancer cells exhibiting Paneth cell properties in humans, we conducted an analysis of scRNA-seq data obtained from colorectal cancer patients (36). Our analysis focused on CD45- EpCAM+ epithelial cells, excluding hematopoietic cells. UMAP plotted 31 clusters of single cells derived from normal tissue, primary tumor and metastasis samples (Fig. 4A and B). Employing lysozyme expression and Paneth cell module scores (Pangalo database) as criteria, we have identified LYZ+ cancer cells harboring Paneth cell properties across 5 clusters (Fig. 4C-H, Fig. S4 and S5). LYZ+ cancer cell population is distinct in metastasis samples (Fig. 4E and 4F). LYZ expression over 2.0 (3^rd^ Quantile, round 1) and Paneth module scores over 0.2 (3^rd^ Quantile, round 2 to avoid getting 0) single cells were identified as LYZ+ cancer cells harboring Paneth cell properties, 1%, 13% and 24% in normal, primary tumor and metastasis samples, respectively.

**Figure 4.**
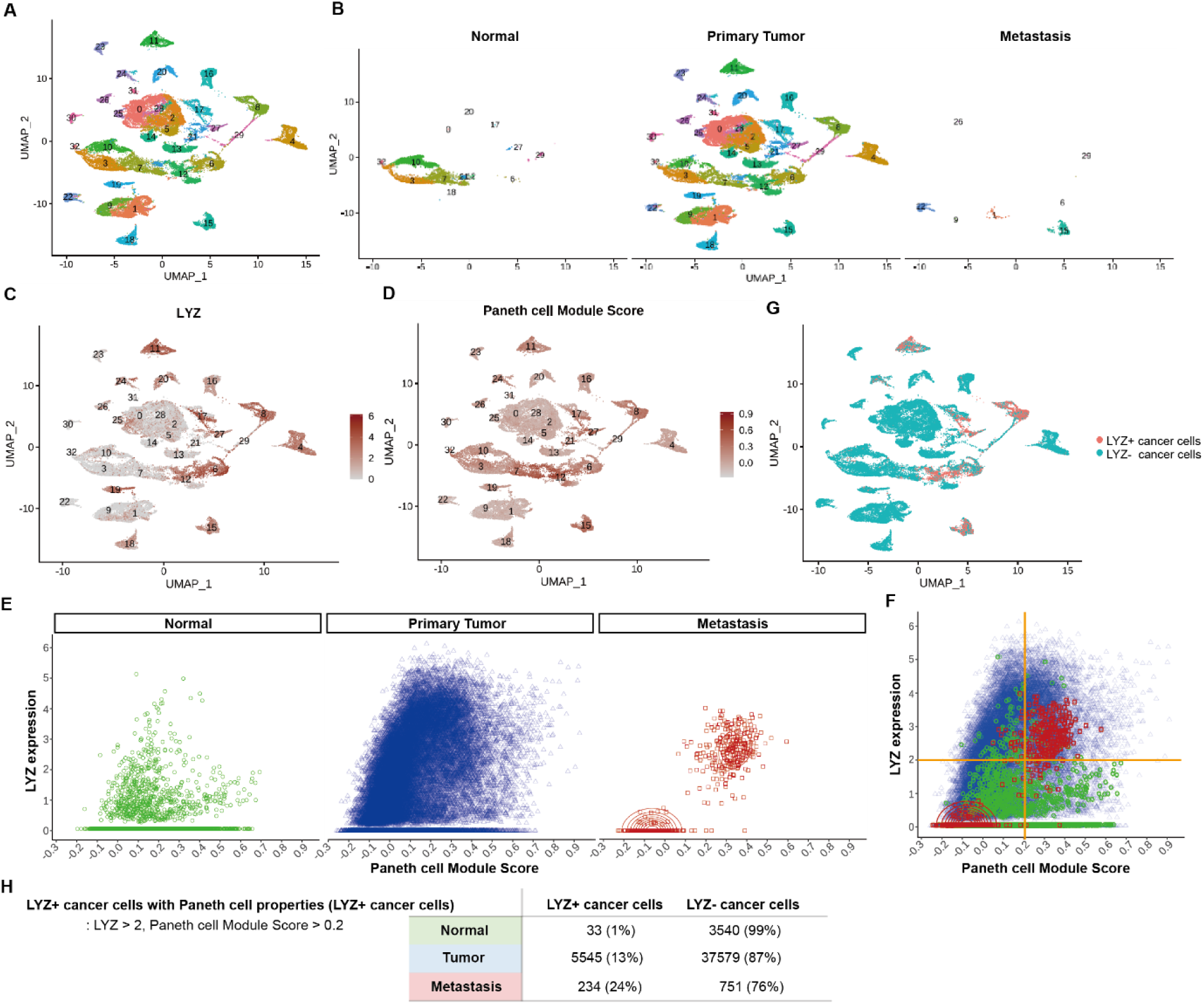
Identification of colon cancer cells harboring Paneth cell properties in humans. Published colorectal cancer patient scRNA-seq data (REF) was analyzed to identify the presence of cancer cells with Paneth cell properties. **(A)** The UMAP plot of total 31 clusters. **(B)** Normal cells, primary tumor cells and liver metastasized cells (Metastasis) are shown in the UMAP plot clusters. **(C-D)** Expression levels of lysozyme (LYZ) and Paneth cell module scores are displayed in the UMPA plot. **(G)** Based on the analysis in (C) and (D), cancer cells harboring Paneth cell properties are indicated as red dots (LYZ+ cancer cells). **(E)** Lysozyme expression and Paneth cell module scores of single cells in Normal, Primary Tumor and Metastasis samples are presented by dot plots. **(F)** Dot plots in (E) are overlayed. **(H)** The percentiles of cancer cells with Paneth cell properties (LYZ+ cancer cells) are shown.

This delineates the distribution of LYZ+ cancer cells with Paneth cell characteristics in different stages of colorectal cancer progression.

To unravel the functional significance of LYZ+ cancer cells, we performed gene set enrichment analysis for the scRNA-seq data obtained from colon cancer patients. In correlation with our murine scRNA-seq data IPA results, glycolysis emerged as the 2^nd^-ranked enriched pathway in LYZ+ cancer cells compared to LYZ- cancer cells (Fig 5A). Notably, key players in glycolysis such as *ALDOB* and *KDELR3* genes are increased in LYZ+ cancer cells (Fig. 5B). Also, the hallmark of protein secretion was the top-ranked enriched pathway, which is essential for Paneth cell activity including secretion of anti-microbial peptides (Fig. 5A). These results suggest the existence of a distinct population of LYZ+ colon cancer cells endowed with Paneth cell properties. Collectively, DKK2 is required for the generation of LYZ+ cells to form the cancer stem cell niche, a prerequisite for the outgrowth of metastasized colorectal cancer cells.

**Figure 5.**
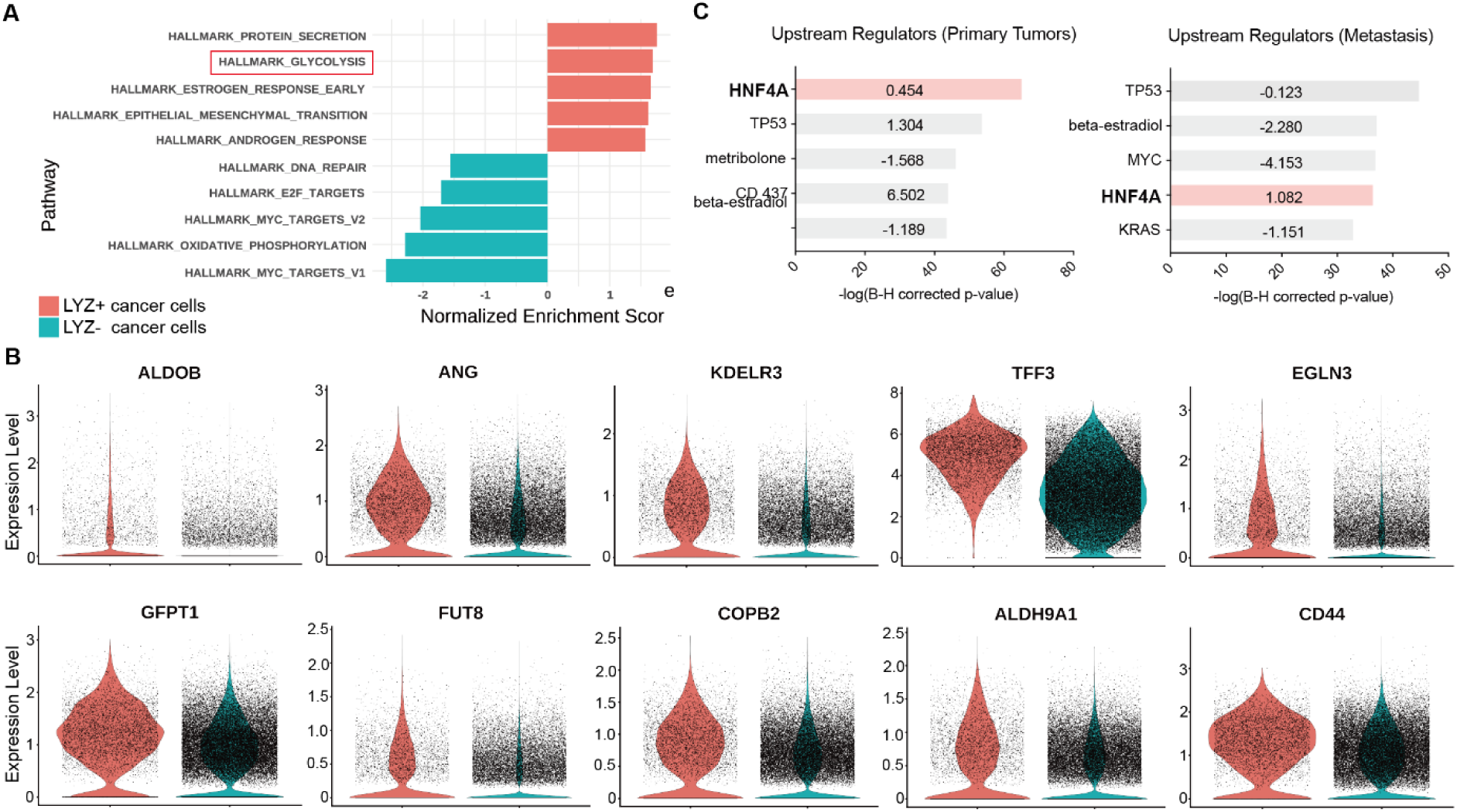
Colon cancer cells with Paneth cell properties contribute to glycolysis. Colorectal cancer patient scRNA-seq data were further analyzed by Gene set enrichment analysis (GSEA) and Ingenuity Pathway Analysis (IPA) **(A)** GSEA of cancer cells with Paneth cell properties (LYZ+ cancer cells) compared to all other cancer cells (LYZ-cancer cells), shown in Fig.4 (G). **(B)** Representative gene expressions in the hallmark of glycolysis pathway are shown. **(C)** Upstream regulators suggested by IPA in Primary Tumor and Metastasis are presented. Activation z-scores are indicated in the bar. IPA predicted activation or inhibition of the upstream regulators are colored as red and cyan, respectively.

### HNF4A mediates the formation of Lysozyme positive colon cancer cells by DKK2

Ingenuity pathway analysis (IPA) of the human scRNA-seq data from primary tumor and metastasis samples suggested hepatocyte nuclear factor 4 alpha (HNF4A) as an upstream regulator of LYZ+ cancer cells compared to LYZ- cancer cells (Fig. 5C). In line with this, IPA of the bulk RNA-seq data using mouse colon cancer organoids suggested that HNF4A is activated in KO organoids (Fig. 6A). This is consistent with our previous report that DKK2 enhanced *Lgr5* expression in colitis-induced cancer cells via HNF4α1, an isoform of HNF4A (28). Significant enrichment of the HNF4A binding motif was also detected in the open chromatin regions of KO organoids compared to AKP controls determined by assay for transposase-associated chromatin using sequencing (ATAC-seq) (Fig. 6B). In comparison with normal colonic organoids, the HNF4A motif was listed as the most reduced motif in AKP indicating significant reduction of HNF4A-regulated transcription whereas it might be rescued by *Dkk2* knockout (Fig. 6C). Notably, HNF4A was the only molecule suggested by all of these multi-omics data analyses in both mouse and human. These results suggest HNF4A as a key transcription factor in downstream signaling of DKK2 in colon cancer cells.

**Figure 6.**
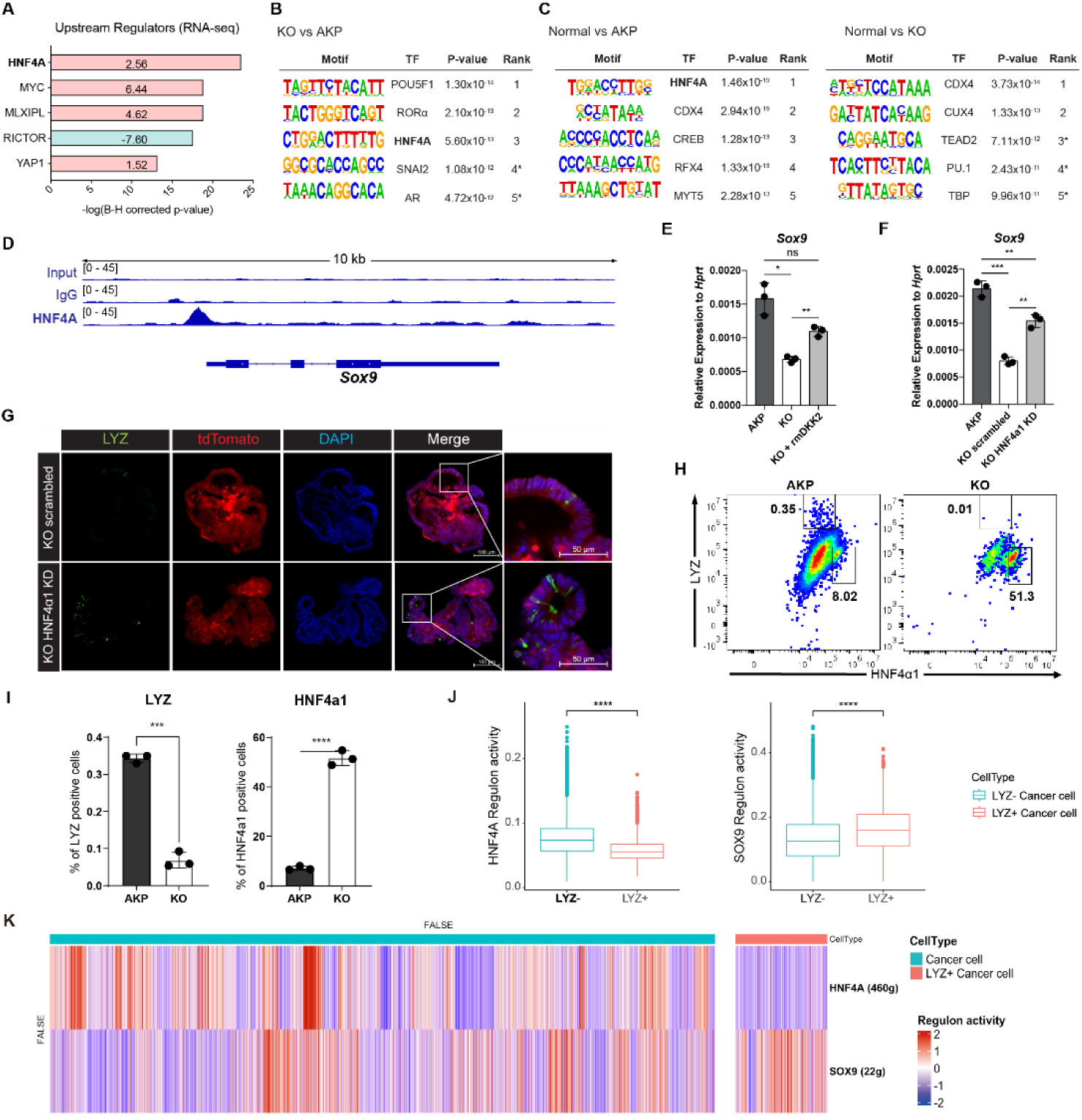
DKK2-driven reduction of HNF4α1 protein in murine colon cancer organoids promotes cancer cells with Paneth cell properties. **(A)** Ingenuity pathway analysis (IPA)-suggested upstream regulators of the RNA-seq data shown in Figure 2A. Z-scores are presented in the bar. IPA predicted activation or inhibition of the upstream regulators are colored as red and cyan, respectively. **(B-C)** List of top 5 transcriptional factors (TF) in the motif enrichment analysis of ATAC-seq data comparing normal colonic organoids, AKP and KO cancer organoids. * indicates possible false positive. **(D)** Chromatin immunoprecipitation sequencing (Chip-seq) analysis of KO organoids using an anti-HNF4A antibody. **(E)** Quantitative expression of *Sox9* in colon cancer organoids in the presence or absence of DKK2 (rmDKK2: recombinant mouse DKK2 protein added in organoid culture). **(F)** Analysis of Sox9 expression after knockdown *HNF4*α*1* in KO colon cancer organoids (KO HNF4α1 KD). ns = not significant, *P<0.05, **P<0.01, ***P<0.001; two-tailed Welch’s t-test. Error bars indicate mean ± s.d. **(G)** Representative images of confocal microscopy analysis of Lyz-stained cancer cells with Paneth cell properties in DKK2 KO HNF4α1 KD organoids. **(H-I)** Control AKP or KO colon cancer organoids were transplanted via splenic vein as described in Figure 1. 3 weeks after transplantation, mice were sacrificed to analyze metastatic tumor growth in liver. Quantification of cancer cells with Paneth cell properties in metastasized tumor nodules by flow cytometry for Lysozyme (LYZ) and HNF4α1. Tumor cells were initially gated by the tdTomato reporter expression. Representative images of flow cytometry are shown (H). Statistic analyses of the percentiles of LYZ positive cells (% of upper left) and HNF4α1 positive cells (% of lower right) in tumor nodules (I). **P<0.01, ****P<0.0001; two-tailed Welch’s t-test. Error bars indicate mean ± s.d. 3 mice were tested per group. Data are representative of two independent experiments. **(J-K)** Reduced HNF4A regulation activity with enhanced SOX9 regulation activity in LYZ+ cancer cells in human colorectal cancer scRNA-seq data. Box plots represent the regulon activity of HNF4A and SOX9 in LYZ+ cancer cells. ****P<0.0001; Wilcoxon signed-rank test (J). Z-scaled regulon activities of HNF4A and SOX9 in human colon cancer cells is displayed by heatmap (K).

HNF4A is necessary for maturation of fetal intestine and barrier functions of intestinal epithelium in the homeostatic condition (37, 38). Two *Hnf4a* isoforms—*Hnf4α1* and *Hnf4α7*— are involved in intestinal homeostasis and regeneration (39). Intestinal inflammation induces expression of the HNF4α1 isoform in the crypt epithelial cells to regenerate the epithelium whereas loss of the HNF4α1 isoform occurs in colitis-induced tumor cells (39). Loss of the HNF4α1 isoform was also observed in AKP organoids while *Dkk2* knockout restored the presence of HNF4α1 protein in the nucleus of AKP organoid cells (28). This allowed us to investigate the binding regions of HNF4α1 in colon cancer cells using KO organoids. Chromatin immunoprecipitation followed by sequencing (Chip-seq) analysis using the Chip-seq grade anti- HNF4A antibody revealed that HNF4A binds to the promoter region of *Sox9* in KO organoid cells (Fig. 6D). Sox9 is a transcription factor required for the secretory lineage of epithelial cell differentiation including Paneth cells (23). Quantitative gene expression analysis showed that *Dkk2* knockout reduced *Sox9* expression in AKP organoids, which was rescued by recombinant DKK2 protein treatment (Fig. 6E). Reduced *Sox9* expression in KO organoids was also reversed by HNF4α1 knockdown (Fig. 6F). These data indicate that DKK2-driven loss of HNF4α1 protein leads to *Sox9* expression followed by formation of LYZ+ colon cancer cells. Confocal microscopy analysis confirmed that HNF4α1 knockdown rescued the presence of LYZ+ cancer cells in KO organoids (Fig. 6G). In order to quantify LYZ+ cancer cells in the mouse model of liver metastasis, tdTomato positive cells in the liver were identified as the implanted organoid-derived cancer cells then, LYZ+ cells were calculated (Fig. 6H). The percentile of LYZ+ cells in total cancer cells was decreased about 3-fold in *Dkk2* knockout metastasized cells (KO) compared to the AKP control cells in liver (Fig. 6I). Simultaneously, HNF4α1 positive cells were tripled to about 60 percent in KO (Fig. 6H and 6I). These findings suggest that DKK2 is indispensable for the generation of LYZ+ cancer cells in liver metastasized nodules by reducing protein levels of HNF4α1. Further analyses for human colorectal cancer scRNA-seq data revealed that HNF4A regulation activity is reduced in LYZ+ human colon cancer cells compared to the LYZ- cancer cells while SOX9 regulation activity is significantly increased (Fig. 6J-6K and S5) suggesting the key roles of HNF4A and SOX9 in LYZ+ cancer cell formation in human. Taken together, DKK2- driven loss of HNF4α1 protein enhances Sox9 expression in colon cancer cells to generate LYZ+ cells with Paneth cell properties.

## Discussion

We have shown that DKK2 is indispensable for metastatic tumor growth of colorectal cancer in the murine model developed by transplantation of colon cancer organoids carrying mutation in the *Apc*, *Kras* and *Tp53* genes. Bulk RNA-seq analysis of DKK2-deficient colon cancer organoids showed reduction of the marker genes of Paneth cells, which play a role in the stem cell niche development. Single cell RNA-seq analyses for mouse and human metastatic colon cancers have identified the existence of LYZ+ cancer cells harboring Paneth cell properties, in particular, glycolysis for stem cells. In the absence of DKK2, both pathway enrichment analysis of the RNA-seq and ATAC-seq data suggested that HNF4A is activated. This is consistent with our previous report in the murine model of colitis-induced tumorigenesis (28). ChIP-seq analysis using an anti-HNF4A antibody revealed that HNF4A directly binds to the promoter region of *Sox9*, a transcriptional factor required for epithelial differentiation into Paneth cells (22, 23). Notably, DKK2 induces the loss of HNF4α1 isoform in colon cancer cells followed by elevation of *Sox9* expression lead to the formation of LYZ+ cancer cells exhibiting Paneth cell properties. DKK2- deficient colon cancer cells possessed high levels of HNF4α1 protein and failed to generate LYZ+ cancer cells *in vitro* and *in vivo*. DKK2 deficiency resulted in reduced glycolysis in mouse liver metastasized colon cancer cells. These results demonstrate the significant reduction of liver metastasis in DKK2 knockout (KO) colon cancer cells infused mice.

*Fumagalli et al.* have recently shown that the generation of *Lgr5* positive cells from the metastasized *Lgr5* negative cancer cells is necessary for their outgrowth using *in vivo* time-lapse imaging and organoid culture of metastatic cancer cells that escaped from the primary colon cancer tissues (7). This finding implies the necessity of both cancer stem cells and their niche for metastatic outgrowth. Indeed, Paneth cell-derived stem cell niche factors such as Wnt3, EGF and Dll4 are essential for the maintenance of stem cells (11). Dietary induced mouse sporadic intestinal cancer showed elevation of Paneth cell marker genes in the intestinal epithelium (40). IDO1+ Paneth cells contribute to the immune evasion of colon cancer (41). Depletion of Paneth cells or Paneth cell-derived Wnt3 in *Apc^Min^*mice impaired intestinal adenoma formation (42). The plasticity of Paneth cells allows dedifferentiation of Paneth cells into intestinal stem cells (15, 43). These reports indicate the necessity of the Paneth cells during the development of colon cancer.

However, the existence of Paneth cells in large intestine or tumor tissues remains unclear. In 2016, Reg4+ secretory cells have been suggested as Paneth-like cells in the colon (24). Single cell transcriptome analysis showed the presence of Paneth-like cells in human colon (25). Recent study further supported the presence of Paneth-like cells within the colonic stem cell niche during bifidobacterium-induced intestinal stem cell regeneration (26). Our study suggests the presence of LYZ+ cells exhibiting Paneth cell properties in the context of colon cancer and metastasis. This extends our knowledge of colorectal cancer progression that DKK2 is required for the generation of LYZ+ cells forming cancer stem cell niches.

Further investigation will be needed to characterize LYZ+ cancer cells with Paneth cell properties observed in colon cancers carrying mutations in *Apc*, *Kras* and *Tp53* genes compared to normal Paneth cells in order to identify their roles in cancer stem cell niche development.

Recent studies have reported stem cell niche remodeling in the primary tumor in the presence of oncogenic mutations. Cancer stem cells outcompete their neighboring normal stem cells during intestinal tumorigenesis, which is known as clonal fixation (44). *Apc*-mutant cancer cells secrete several Wnt antagonists such as NOTUM to inhibit proliferation of normal intestinal stem cells and facilitate their differentiation (45). Other oncogenic mutations in *Kras* or *Pi3kca* genes lead to paracrine secretion of BMP ligands, remodeling the stem cell niche that is detrimental for maintaining normal stem cells (46). These changes are beneficial for the outgrowth of cancer stem cells in the crypt where the stem cell niche had been developed prior to tumorigenesis. In the context of metastasis, the formation of stem cell niches is required upon metastases seeding in the liver. We have shown that DKK2 is required for the formation of LYZ+ cancer cells carrying Paneth cell properties. DKK2 expression is induced by *Apc* knockout and further increased by additional oncogenic mutations in *Kras* and *Tp53* genes (28). Characterization of DKK2-induced LYZ+ cancer cells will provide better understanding of the biology of cancer stem cell niches, particularly in liver metastases of colorectal cancer.

DKK2 expression in colon cancer promotes LYZ+ cancer cell generation through the regulation of HNF4α1, which suppresses expression of *Sox9,* the transcription factor for Paneth cell differentiation. It has been recently reported that HNF4α2 isoform acts as an upstream regulator of Wnt3 and Paneth cell differentiation implying the roles of HNF4A for Paneth cell differentiation in the homeostatic condition (47). Using *Villin^Cre^-Apc^floxed^* mice, Suzuki et al. have confirmed that HNF4A proteins are completely absent in cancer cells with high β-catenin activity along with increased *Sox9* expression (48). *Sox9* regulates cell proliferation and is required for Paneth cell differentiation (22, 23). Bi-allelic inactivation of the *Apc* gene induces *Sox9* expression in colon cancer (49). Likewise, *Apc* inactivation induces *Dkk2* expression (28). Here we showed that DKK2-mediated HNF4α1 protein degradation enhanced *Sox9* expression in colon cancer (Fig. 7). Notably, *Apc* mutation-driven *Sox9* expression blocks intestinal differentiation and activates stem cell-like program (49). This is consistent with our previous report that DKK2 enhances *Lgr5* expression in colon cancer (28). In sum, our findings suggest that a Wnt ligand, DKK2 is an upstream regulator of SOX9 expression for enhanced stem cell activity via the formation of LYZ+ cells with Paneth cell characteristics in colorectal cancers.

**Figure 7.**
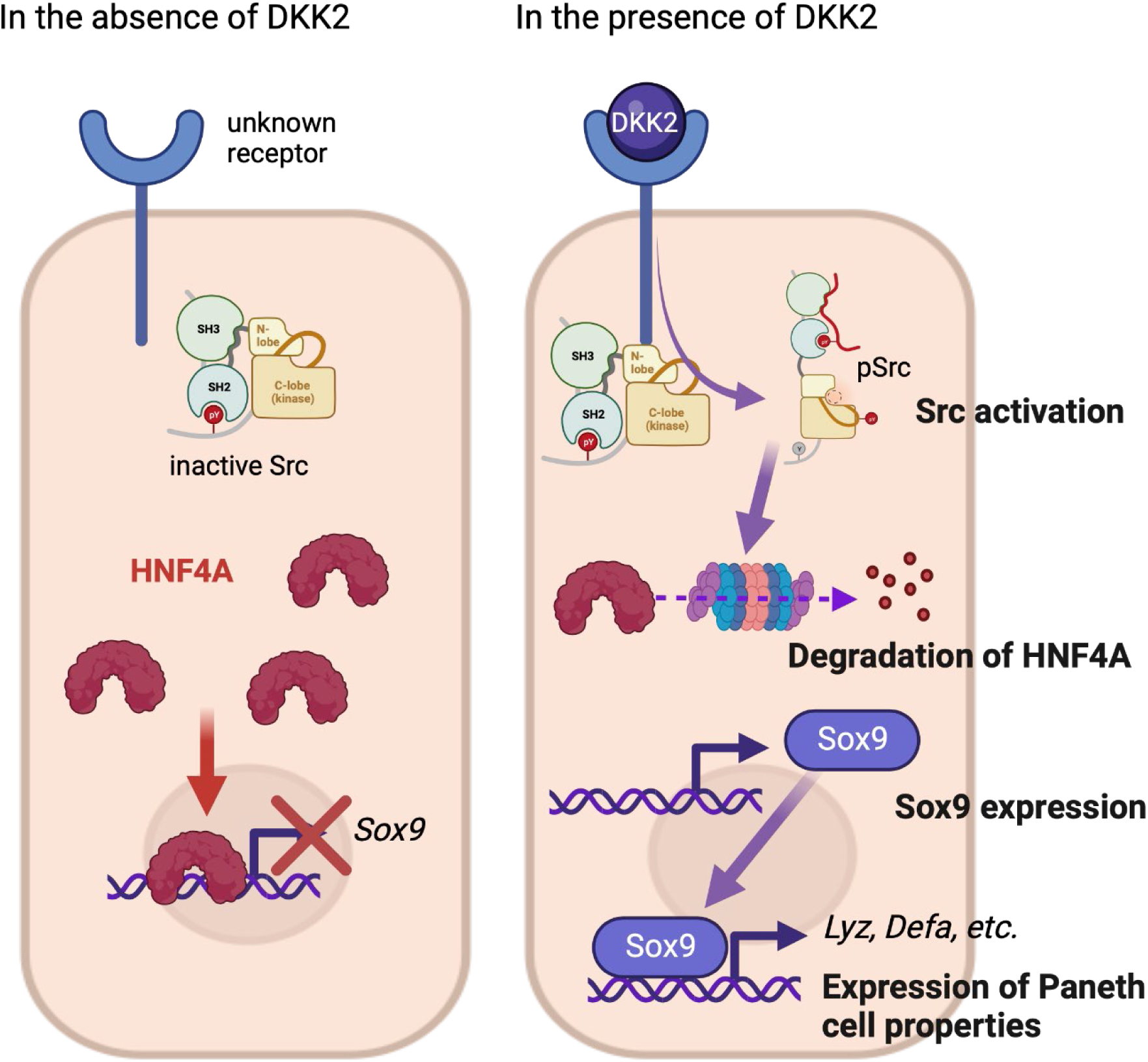
The suggested mechanism of DKK2 in the formation of colon cancer cells with Paneth cell properties. In the absence of DKK2, Sox9 expression is inhibited by HNF4A. Our previous report has shown that DKK2 activates Src followed by degradation of HNF4A protein (28). HNF4A deficiency leads to Sox9 expression in colon cancer cells that induces Paneth cell properties including the expression of lysozymes and defensins. Formation of cancer cells with Paneth cell properties by DKK2 contributes the outgrowth of metastasized colon cancer cells in the liver.

In this study, we have shown that DKK2 promotes liver metastasis of colon cancer. DKK2 is required for the formation of LYZ+ colon cancer cells exhibiting Paneth cell properties such as glycolysis for stem cells. Glycolysis in *Lgr5* expressing cancer cells was reduced along with the loss of LYZ+ cells by *Dkk2* knockout. As a result, metastatic growth of colon cancer cells was significantly inhibited *in vivo*. These findings suggest the necessity of DKK2 in the development of cancer stem cell niches for developing metastases. Mechanistically, DKK2-mediated degradation of HNF4α1 protein elevated *Sox9* expression followed by LYZ+ cell formation. Given that DKK2 expression is intestinal tumor specific, inhibition of DKK2 may have promising therapeutic outcomes in the treatment of metastatic colon cancer.

## Materials and Methods

### Splenic injection

Generation of AKP, *Dkk2* knockout AKP, *HNF4*α*1* knockdown and *Dkk2* knockout AKP organoids and their culture protocols have been published in the previous study (28). We used the previously reported splenic injection methodology to develop organoid-derived liver metastasis in mice (50). Cultured organoids were prepared as single cells following 10-30 min of digestion in TrypLE solution (Thermo, 12605010) at 37°C, followed by centrifugation with advanced DMEM- F12 (Thermo, 12634010). The isolated single cells were re-suspended with PBS as 300,000 cells per 100 μl of PBS. Surgical instruments were sterilized by steam autoclave (250 ° F., 15 p.s.i. for 30 minutes). 8-12 weeks old C57BL/6 mice were anesthetized with Ketamine (100 mg/kg) plus Xylazine (10 mg/kg) via intraperitoneal injection. Puralube Vet ointment was applied to the eyes to prevent drying while surgery. Spleen was divided by two Horizon medium size ligating clips (Teleflex, 002200) in the center of the spleen and one hemi-spleen was returned into the peritoneum. Using a 26-½ Gauge needle, 100-200 μl volume of cells was injected underneath of the splenic capsule over the course of 30 s. A small Horizon clip (Teleflex, 001200) was applied to ligate the pancreas and splenic vessels and the injected hemi-spleen was removed. The incision was closed with a 5-0 running stitch using Vicryl (Ethicon, J493G) and 2-3 skin clips. Meloxicam (0.3 mg/kg) was injected subcutaneously to alleviate postsurgical pain at 24-72 h post-surgery.

### In vivo imaging

Noninvasive *in vivo* imaging was performed using a Microfocus X-ray imaging source (Kodak). Mice were anesthetized by isoflurane inhalation and their abdominal hair was shaved. Tandem repeat Tomato (tdTomato) fluorescence in liver metastatic tumors was recorded for 30 sec. tdTomato signals were statistically analyzed using the Bruker image analysis system. X-ray image was captured by 10 sec exposure and merged with the tdTomato image.

## Data availability

All data are included in the article and SI Appendix. All the sequencing data reported in this study have been deposited in the Gene Expression Omnibus as follows; Bulk RNA-seq: GSE157531, ATAC-seq: GSE157529, Chip-seq: GSE277510, scRNA-seq: GSE157645.

## Supporting information

Responses to Reviewers

## Acknowledgements

The authors would like to thank Gouzel Tokmulina for the FACS sorting.

## Author contributions

J.H.S. and A.L.M.B. designed research; J.H.S., J.L., J.J., R.K.D, S.E.M., and J.Y.H. performed research; J.H.S., J.P., J.L., J.J., R.K.D., S.E.M., J.Y.H., and A.L.M.B. contributed new reagents/analytic tools; J.H.S., J.P., J.J., J.L., J.C., J.W. and A.L.M.B. analyzed data; J.H.S., J.P., J.K., S.P., J.C. performed the revision experiments; and J.H.S., J.J., J.L., J.C. and A.L.M.B. wrote the paper.

This work was supported by NIH RO1 CA168670-01 (awarded to A.L.M.B), Yonsei University Future-Leading Research Initiative 2023-22-0438 (awarded to J.H.S.) and National Research Foundation of Korea (NRF) RS-2023-00213586 (awarded to J.H.S.).

