## Supplemental Information for "Metastasis of colon cancer requires Dickkopf-2 to generate cancer cells with Paneth cell properties"

### Supplementary figures and table

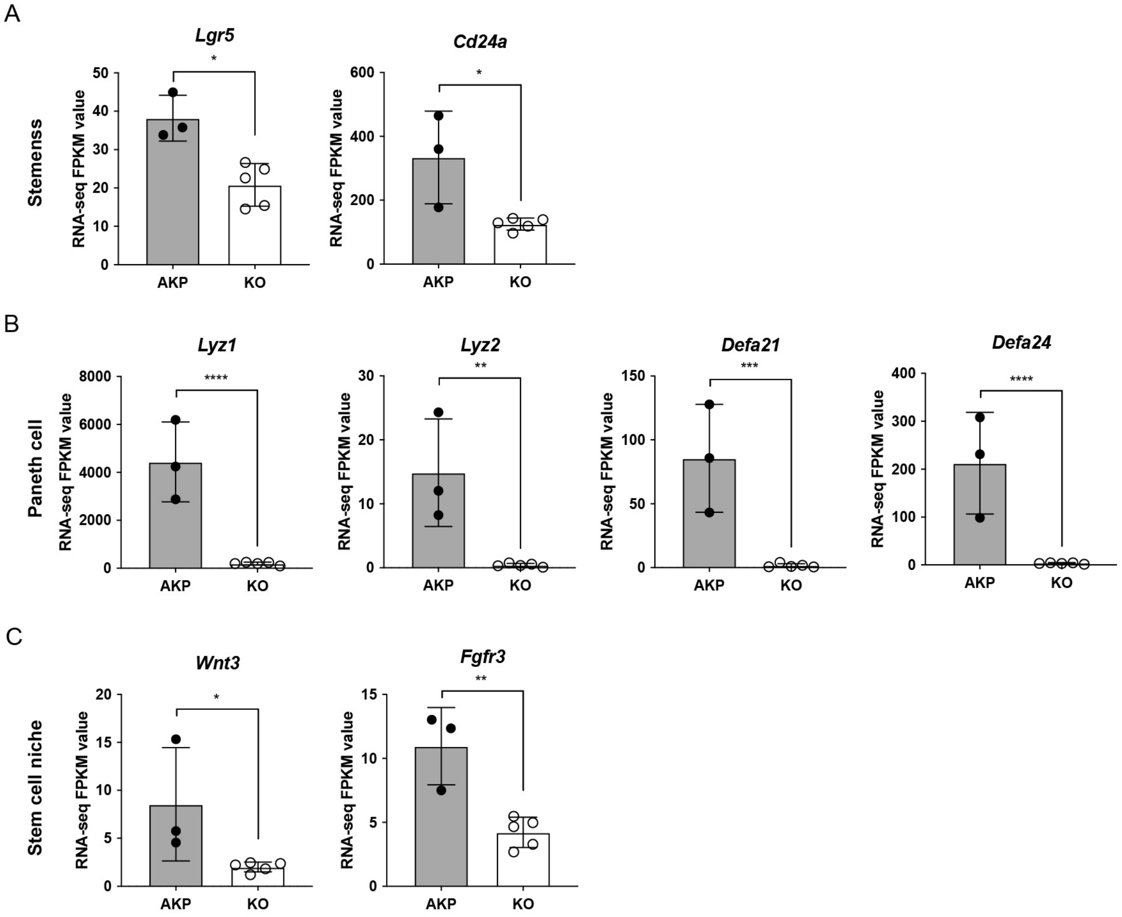


**Figure S1. Bulk RNA sequencing (RNA-seq) analysis of *Dkk2* knockout cancer organoids.**

**(A-C)** Bulk RNA-seq analysis was performed using 8 days cultured colon cancer organoids carrying mutations in *Apc*, *Kras* and *Tp53* genes. Expression of the marker genes of stemness (A) and Paneth cells (B) are shown. *Cd24a* is a marker of both stemness and Paneth cells. Expression of genes related in the stem cell niche is shown in (C). *P<0.01, **P<0.05, ***P<0.01, ****P<0.001; adjusted p-values (False Discovery Rate) by Sleuth (see Supplementary Methods). 3 and 5 biological replicates of control (AKP) and *Dkk2* knockout (KO) cancer organoids were tested, respectively.

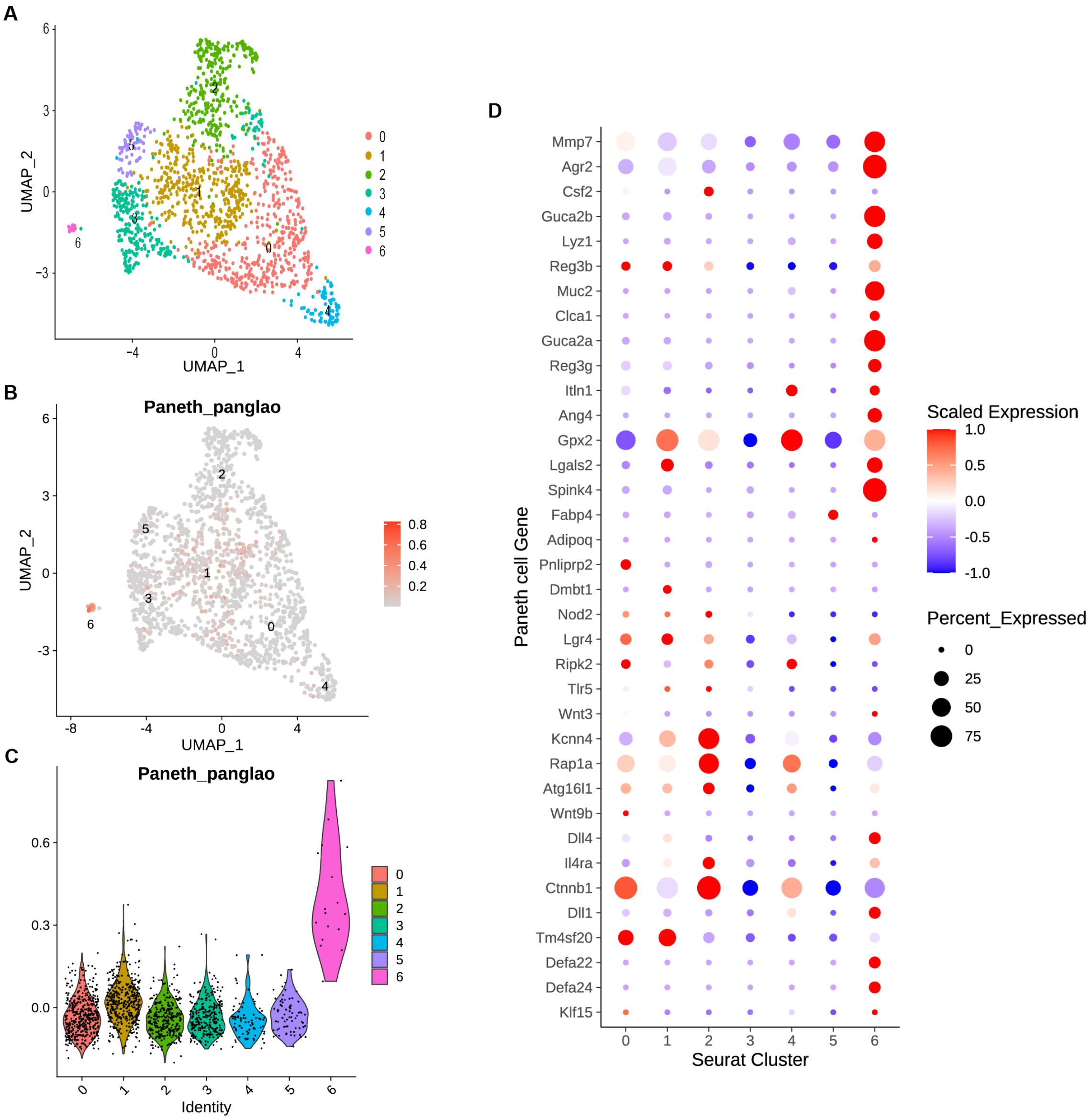


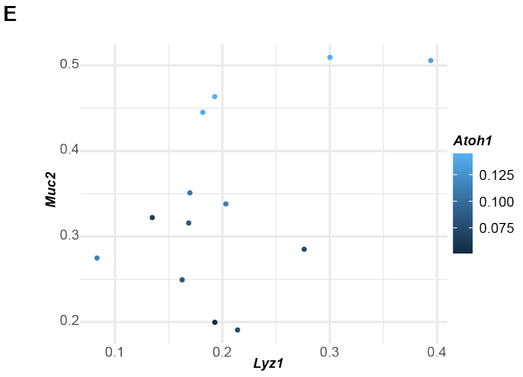


F

| | AKP | KO |
|---|---|---|
| Paneth+ cells (*Lyz1+Muc2+*)/cluster 6 | 2/6 | 3/11 |
| Goblet+ cells (*Muc2+Atoh1+*)/cluster 6 | 2/6 | 7/11 |
| Paneth+ cell/Goblet+ cell ratio | 1 | 0.43 |

**Figure S2. Identification of the cluster of Paneth-like cells in the scRNA-seq data of liver metastasized murine colon cancer cells.**

**(A)** The UMAP plot of tumor epithelial cells revealed 7 clusters. **(B)** Paneth cell module score is displayed in the UMAP plot. **(C)** The violin plot displays the Paneth cell module score. It is the highest in the cluster 6. **(D)** A gene expression dot plot with Paneth cell marker genes in epithelial cell clusters. **(E)** A scatter plot of *Lyz1*, *Muc2* and *Atoh1* expression in cluster 6 cells. **(F)** The ratio between cancer cells with Paneth cell properties (Paneth+) and cancer cells with goblet cell properties (Goblet+) in AKP and KO scRNA-seq data is shown.

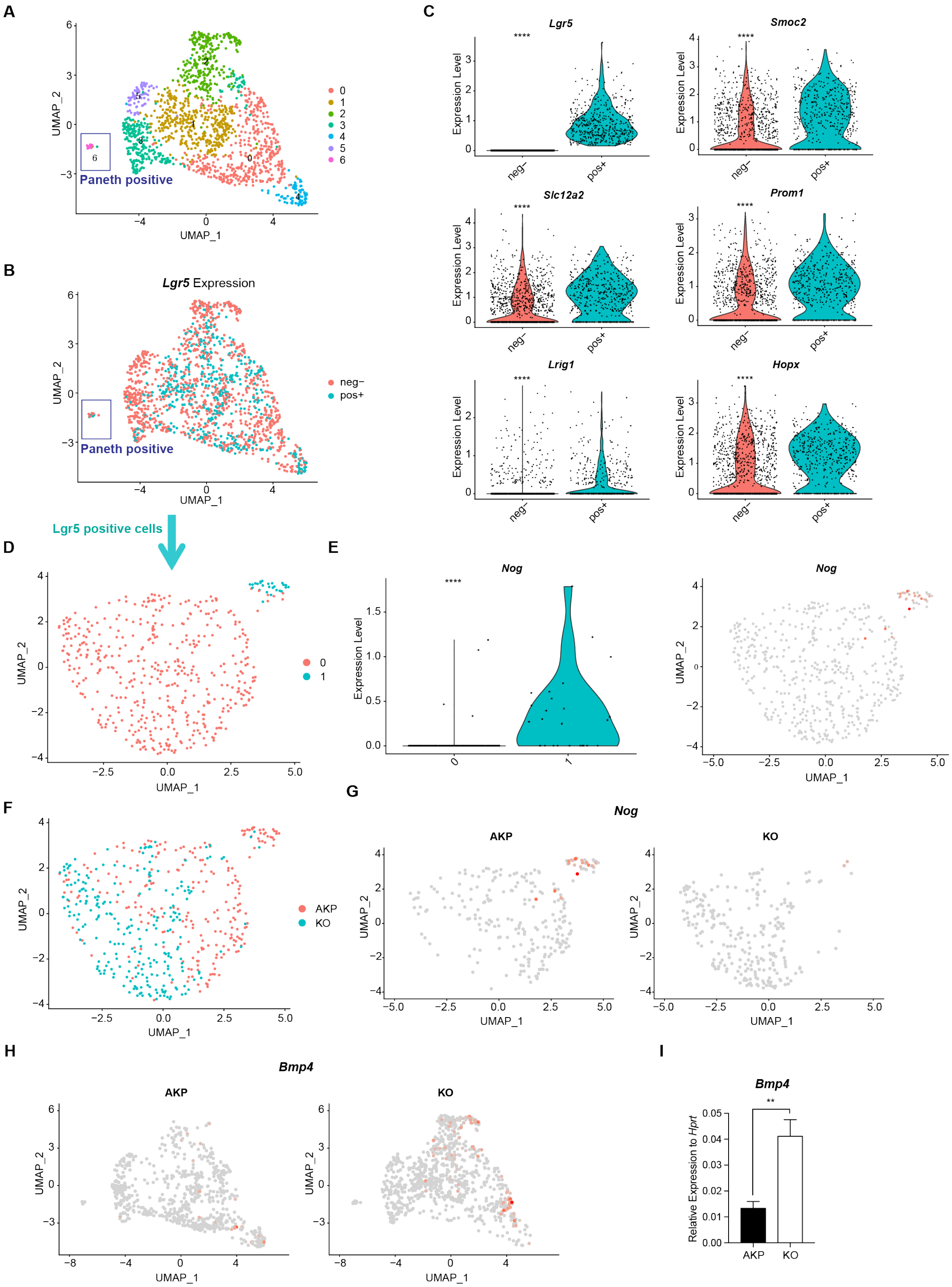


**Figure S3. Reduced expression of *Noggin (Nog)* and reversed expression of *Bmp4* in the scRNA-seq of *Dkk2* knockout metastasized cancer cells.**

**(A)** The UMAP plot of tumor epithelial cells as described in Figure 4C. **(B)** Tumor epithelial cells are separated by *Lgr5* expression (positive: pos+, negative: neg-). **(C)** Differentially expressed genes between *Lgr5* positive and negative cells are presented by violin plots. ****P<0.0001; Wilcoxon signed-rank test. **(D)** The UMAP plot of *Lgr5* positive cells revealed 2 clusters. **(E)** A violin plot displays *Noggin (Nog)* expression mostly detected in the cluster 1 of *Lgr5* positive cells in the UMAP plot showed in (D). ****P<0.0001; Wilcoxon signed-rank test. **(F)** UMAP plot of *Lgr5* positive cells in AKP and KO colored by red and cyan, respectively. **(G)** Expression of *Nog* in the UMAP plots of AKP and KO in (F). **(H)** UMAP plots of *Bmp4* expression in total tumor epithelial cells of AKP and KO in the scRNA-seq data. **(I)** Quantitative PCR analysis of *Bmp4* expression in 8 days cultured colon cancer organoids. **P<0.01; two-tailed Welch's t-test. Error bars indicate mean ± s.d. three biological replicates per group were tested.

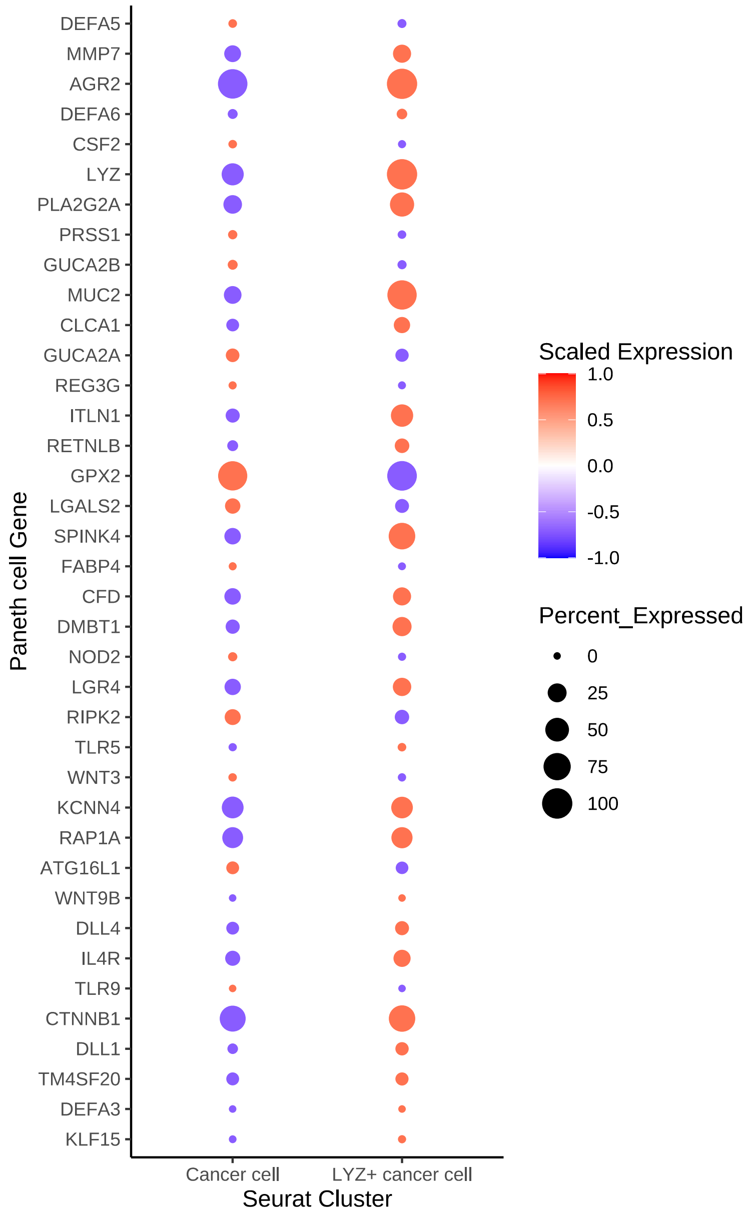


**Figure S4. Paneth cell markers expression in colorectal cancer patients scRNA-seq data.**

A dot plot of Paneth cell marker genes in scRNA-seq data obtained from colorectal cancer patients. Paneth cell marker genes were retrieved from the Panglao database (PangaloDB).

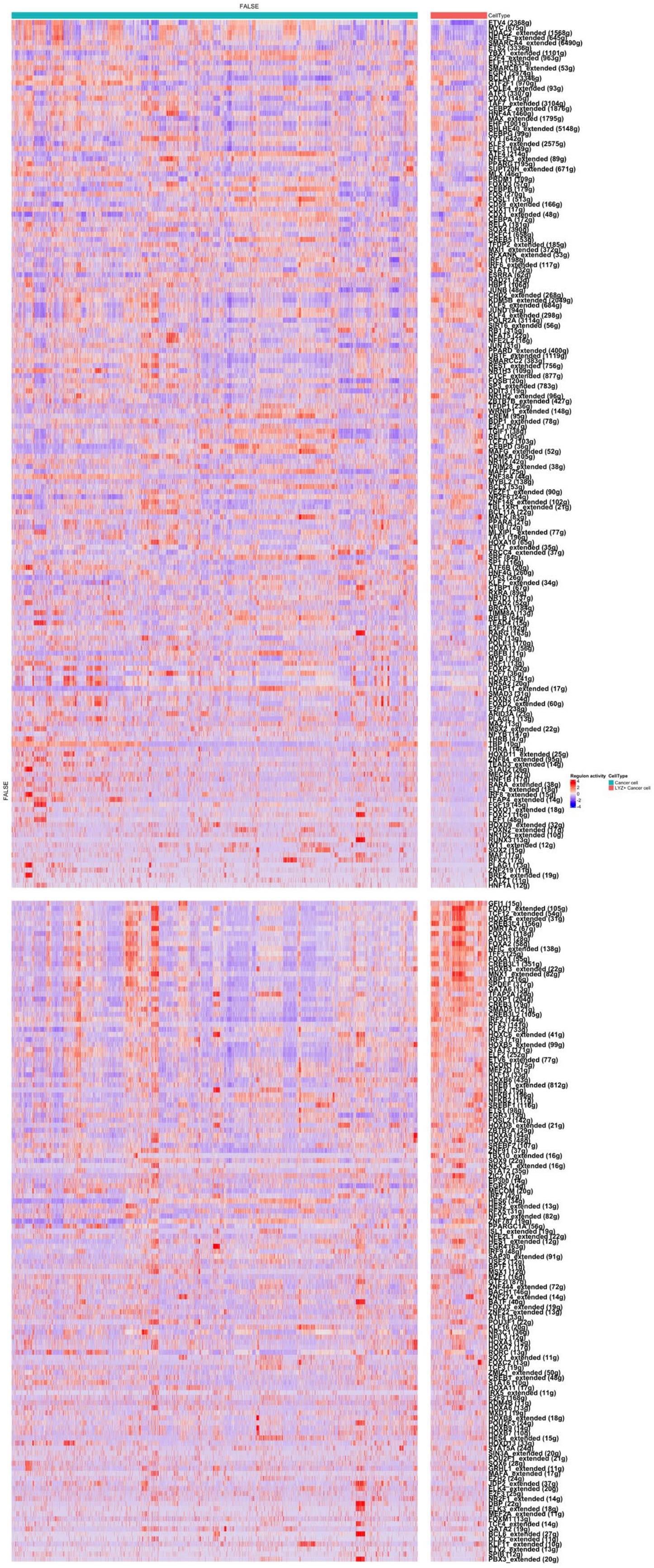
FALSE
CellType
FALSE
Regulon activity
CellType
Cancer cell
LYZ+ Cancer cell

**Figure S5. Analyses of the regulon activity of various transcription factors in lysozyme positive (LYZ+) colon cancer cells in human colon cancer scRNA-seq data.**

The z-scaled regulon activities of various transcription factors in LYZ+ cancer cells compared to LYZ- cancer cells are displayed by heatmap. The regulon activity is calculated based on the expression of transcription factors and their downstream target molecules.

**Table S1. The list of primers used in quantitative real time PCR**

| | | |
|---|---|---|
| *Dkk2* | Forward (5' -3') | GTACCCGCTGCAATAATGGAATCT |
| | Reverse (5' -3') | AACAGACTCAGCACAGCGAA |
| *Hnf4α1* | Forward (5' -3') | ATGCGACTCTCTAAAACCCTTG |
| | Reverse (5' -3') | ACCTTCAGATGGGGACGTGT |
| *Hprt* | Forward (5' -3') | CTCCTCAGACCGCTTTTTGC |
| | Reverse (5' -3') | TCATCGCTAATCACGACGCT |
| *Lgr5* | Forward (5' -3') | AGCCTATGGACTCAATGTGAAGA |
| | Reverse (5' -3') | AAATCAGCCCTAGGTCAAGATGATA |
| *Lyz1* | Forward (5' -3') | GAGACCGAAGCACCGACTATG |
| | Reverse (5' -3') | CGGTTTTGACATTGTGTTCGC |
| *Lyz2* | Forward (5' -3') | ATGGAATGGCTGGCTACTATGG |
| | Reverse (5' -3') | ACCAGTATCGGCTATTGATCTGA |
| *Sox9* | Forward (5' -3') | CGGAACAGACTCACATCTCTCC |
| | Reverse (5' -3') | GCTTGCACGTCGGTTTTGG |

### Supplementary Materials and Methods

#### *Tumor dissociation*

Liver metastasized tumors were collected and dissociated in the digestion buffer containing 200 U/ml of type IV collagenase (Worthington, LS004188), 125 μg/ml of type II dispase (Sigma, D4693), 2.5 % fetal bovine serum, 1x Penicillin/Streptomycin in DMEM at 37 °C for 30 min with agitation. Isolated cells were centrifuged at 400 g and 4 °C for 5 min, then re-suspended in PBS and filtered through the 40 μm pore size strainer.

***Flow cytometry***

Dissociated metastatic tumor cells were fixed and permeabilized with the intracellular fixation and permeabilization buffer set (eBioscience, 88-8824-00) following the manufacturer's protocols. Those cells were stained with DyLight488 conjugated anti-Lgr5 antibody (Clone: OTIA2, Origene, TA400002), anti-phospho-Src (Ty416) antibody (Clone: 9A6, Milliporesigma, 05-677) and Brilliant violet 421 (BV421) Rat anti-mouse IgG3 antibody (Clone: R40-82, BD Horizon, 565808) in Figure 1E. In Figure 4A, tumor cells were stained with anti-Lysozyme antibody (Clone: Poly28600, BioLgened, 8600001), APC-conjugated goat anti-rabbit-IgG polyclonal antibody (Invitrogen, A10931), anti-HNF4a1 antibody (Clone: K9218, R&D Systems, PP-K9218-00) and Alexa Fluor 488-conjugated goat anti-mouse IgG2a secondary antibody (Thermo Fisher Scienctific, AB_2535771). Flow cytometry was performed using the Stratedigm-13.

***RNA isolation***

Total RNA of organoids and liver metastasized tissues were extracted using Trizol reagent (Thermo, 15596018). Organoids in Matrigel culture were mechanically disrupted and spun at 400 g and 4°C for 5 min. Pellets were re-suspended with PBS, then dissolved in Trizol. RNA was isolated using miRNeasy kit (Qiagen, 217004) with on-column DNase digestion (Qiagen, 79254) according to the manufacturer's instructions.

***Quantitative real time PCR***

The SMARTer cDNA synthesis kit (Clontech, 634925) was used for cDNA synthesis from total RNA following the manufacturer's protocol. Quantitative PCR was performed using the iTag universal SYBR Green supermix (BIO-RAD, 1725121) on the CFX96 Touch$^{TM}$ real-time PCR detection system. The list of PCR primers is presented in the supplementary table 1.

***RNA-seq analysis***

RNA was purified from colon cancer organoids using QIAGEN miRNeasy kit (217004) with on-column DNase digestion according to the manufacturer's instructions. RNA-seq libraries were constructed following Illumina TruSeq Stranded mRNA protocol (20020594). The RNA-seq libraries were sequenced on the Illumina Nextseq 500 (42 bp paired-end run) and Illumina HiSeq 2500 instrument platform (76 bp single-end sequencing) for organoids and colitis-induced cancer cells, respectively.

The sequences were basecalled by Illumina RTA embedded in NextSeq Control Software by the standard workflow. The sequencing reads were aligned onto Mus musculus GRCm38/mm10 reference genome using the TopHat v2.1.0 software or Kallisto v0.45.0. The mapped reads were transformed into the count matrix with default parameters using the HTSeq v0.8.0 software, then normalized using the DESeq v2 software. Differentially expressed genes (DEGs) were identified using the same software based on a negative binomial generalized linear model. Sleuth was used to analyze statistical significance of kallisto data.

***ATAC-seq***

Eight days cultured organoids were isolated as single cells using TrypLE (Thermo, 12605010) and fluorescent activated cell sorting (FACS). Dead cells were excluded by Zombie aquaTM (Biolegend 423102) staining. Genomic DNAs were extracted by DNA Clean & Concentrator (Zymo). ATAC-seq libraries were constructed with 50K cells from each condition following Omni-ATAC protocol (Illumina 20034197(Corces et al., 2017))

Sequenced reads were trimmed with adaptor sequences (cutadapt v1.9.1, (Martin, 2011)) and mapped to the mouse genome (GRCm38, emsembl release 93) by Bowtie2 (v2.3.4.1, (Langmead and Salzberg, 2012)). Mitochondrial and duplicated reads were removed by SAMtools (v1.9, (Li et al., 2009)) and Picard (v2.9.0, https://broadinstitute.github.io/picard/), respectively. Peaks were found by MACS2 (v.2.1.1, (Zhang et al., 2008)) and visualized by deepTools (v3.1.1, Ramirez et al., 2014, NAR).

***Chip-seq***

Eight days cultured organoids were isolated as single cells using TrypLE (Thermo, 12605010) and washed with PBS three times. Chromatin Immunoprecipitation was performed using the iDeal

ChIP-seq kit for Transcription Factors (Diagenode, C01010054) and recombinant Anti-HNF4-alpha antibody, ChIP Grade (Abcam, ab181604) following the manufacturers' protocols. ChIP-seq libraries were constructed by Yale Center for Genome Analysis (YCGA) using the KAPA HyperPrep kit and sequenced on the Illumina HiSeq 2500 (150 bp paired-end run). Sequenced reads were trimmed with adaptor sequences (cutadapt v1.9.1) and mapped to the mouse genome (GRCm38, emsembl release 93) by Bowtie2 (v2.3.4.1). Mitochondrial and duplicated reads were removed by SAMtools (v1.9) and Picard (v2.9.0), respectively. Peaks were found by MACS3 (v3.0.0a6) and visualized by deepTools (v3.1.1).

***Confocal microscopy***

Organoids were grown in the Nunc™ Lab-Tek™ II Chamber Slide™ System (Thermo Scienctific, 154534PK). Organoids were fixed with freshly prepared 4 % PFA in the PME buffer (50 mM PIPES, 2.5 mM MgCl2, 5 mM of EDTA) for 20 min at room temperature. Fixed organoids were permeabilized with PBS containing 0.5 % Triton X-100. Fixed and permeabilized organoids were washed with PBS containing 0.05 % Tween 20 then blocking was performed using the wash buffer containing 1 % BSA. Organoids were stained with anti-Lysozyme antibody (Clone: Poly28600, BioLgened, 8600001), Alexa488-conjugated goat anti-rabbit-IgG polyclonal antibody (Invitrogen, A32731) and DAPI (Sigma, D9542). Pictures were captured using a Zeiss Axio Observer Z.1 microscope or a Zeiss LSM780 confocal laser-scanning microscope.

***Mouse single-cell RNA data Processing***

scRNA-seq library was generated using Chromium Single Cell 3’ Reagent Kits v3 (10x Genomics) following the manufacturer’s protocol. Biological replicates were used to generate the replicate library with another methodology as described previously. Fastq files were mapped to pre-built mouse reference set (mm10) and were converted to counts using the pipeline “cellranger count”. Cells exhibiting percent.mt > 30%, nFeatures < 200 and nFeatures > 8000 were filtered out. The count matrix was log-normalized with a pseudo-count of 1 (‘NormalizeData’). The features were then scaled and centered (‘ScaleData’). PCA analysis was performed on scaled matrix of 2000 highly variable genes (HVG, 'FindVariableFeatures') with the top 20 principal components (PCs) detected by knee-point. Clustering was performed by first constructing shared nearest-neighbor graph (SNN) and then applying Louvain with resolution of 0.5 ('FindNeighbors' and 'FindClusters').

***Mouse Cell Type Annotation***

Clusters were annotated based on expression of canonical marker genes, including *Ptprc* for immune cells, *Epcam*, *Krt7*, and *Klf5* for Tumor cells, *Alb*, *Ttr*, and *Crp* for hepatocytes, *Pecam1*, *Eng*, and *Cd34* for Endothelial cells.

Tumor cells were than subsetted, subjected to scaling, PCA analysis on 2000 HVG scaled data with the top 20 PCs, and clustering with resolution of 0.4. Among 6 clusters, cluster 6 exhibited high expression of *Lyz1*, *Muc2*, and *Atoh1* which indicated its Paneth cell and Goblet cell properties.

For cancer stem cell niche analysis, we selected tumor cells with *Lgr5* expression > 0. The subsetted dataset were subjected to scaling, PCA analysis on 2000 HVG scaled data with the top 20 PCs, and clustering with resolution of 0.18.

***Human single-cell data Processing***

We retrieved public human Colorectal cancer single-cell data from ref and utilized cells pre-annotated as Epithelial cells. Cells exhibiting percent.mt > 25% and nFeatures < 200 were filtered out. Additionally, we removed cells with PTPRC expression to make sure our Epithelial cells lacks immune cells, yielding 47,682 cells. The count matrix was log-normalized with a pseudo-count of 1 ('NormalizeData'). The features were then scaled and centered ('ScaleData'). PCA analysis was performed on scaled matrix of 2000 highly variable genes (HVG, 'FindVariableFeatures') with the top 20 principal components (PCs) detected by knee-point. Clustering was performed by first constructing shared nearest-neighbor graph (SNN) and then applying Louvain with resolution of 0.5 ('FindNeighbors' and 'FindClusters').

#### *Human Regulon Analysis*

To confirm reduced activity of HNF4A and elevated activity of SOX9 in LYZ+ human colon cancer cells compared to the LYZ- cancer cells, we utilized single–cell regulatory network inference (SCENIC)(6) and obtained a regulon–activity matrix for each cell. Wilcoxon signed–ranked test with p-values adjusted through Bonferroni correction was performed to compare the activity of HNF4A and SOX9 regulon. Further the regulon-activity matrix was z-scaled to generate the heatmap in Figure 7B.

#### *Gene Set Enrichment Analysis*

To characterize LYZ+ human colon cancer cells, gene enrichment analysis (GSEA) using the fGSEA package on HALLMARK canonical pathways in MSigDB v 7.1 (7). GSEA was performed on pre–ranked genes using 10,000 permutations. Gene ranks were calculated based on MAST differential expression result applying the following formula:

$$stat = avg\_log2FC/abs(\frac{avg\_log2FC}{qnorm(p\_val)})$$

For genes with an infinite value of statistics due to a p–value equal to 0 or 1, the statistics was replaced with the highest or lowest value of the statistical metric.

#### *Paneth-cell Module Score Calculation*

We computed Paneth-cell module score for each cell with marker genes in PanglaoDB (8)by implementing ‘AddModuleScore’ function in Seurat (9). For human dataset, we filtered out marker genes used in mouse, and for mouse dataset, we filtered out marker genes for human.
